# Comprehensive analysis of horizontal gene transfer among multidrug-resistant bacterial pathogens in a single hospital

**DOI:** 10.1101/844449

**Authors:** Daniel R. Evans, Marissa P. Griffith, Mustapha M. Mustapha, Jane W. Marsh, Alexander J. Sundermann, Vaughn S. Cooper, Lee H. Harrison, Daria Van Tyne

## Abstract

Multidrug-resistant bacterial pathogens pose a serious public health threat, especially in hospital settings. Horizontal gene transfer (HGT) of mobile genetic elements (MGEs) contributes to this threat by facilitating the rapid spread of genes conferring antibiotic resistance, enhanced virulence, and environmental persistence between nosocomial pathogens. Despite recent advances in microbial genomics, studies of HGT in hospital settings remain limited in scope. The objective of this study was to identify and track the movement of MGEs within a single hospital system using unbiased methods. We screened the genomes of 2,173 bacterial isolates from healthcare-associated infections collected over an 18-month time period to identify nucleotide regions that were identical in the genomes of bacteria belonging to distinct genera. These putative MGEs were found in 196 isolates belonging to 11 different genera; they grouped into 51 clusters of related elements, and they were most often shared between related genera. To resolve the genomic locations of the most prevalent MGEs, we performed long-read sequencing on a subset of representative isolates and generated highly contiguous, hybrid-assembled genomes. Many of these genomes contained plasmids and chromosomal elements encoding one or more of the MGEs we identified, which were often arranged in a mosaic fashion. We then tracked the appearance of ten MGE-bearing plasmids in all 2,173 genomes, and found evidence supporting the transfer of plasmids between patients independent from bacterial transmission. Finally, we identified two instances of likely plasmid transfer across genera within individual patients. In one instance, the plasmid appeared to have subsequently transferred to a second patient. By surveying a large number of bacterial genomes sampled from infections at a single hospital in a systematic and unbiased manner, we were able to track the independent transfer of MGEs over time. This work expands our understanding of HGT in healthcare settings, and can inform efforts to limit the spread of drug-resistant pathogens in hospitals.

## INTRODUCTION

Horizontal gene transfer (HGT) is a driving force behind the multidrug-resistance and heightened virulence of healthcare-associated bacterial infections^1^. Genes conferring antibiotic resistance, heightened virulence, and environmental persistence are often encoded on mobile genetic elements (MGEs), which can be readily shared between bacterial pathogens via HGT^2^. While rates of HGT are not well quantified in clinical settings, prior studies have shown that MGEs can mediate and/or exacerbate nosocomial outbreaks^3–6^. Recent studies have also demonstrated that multidrug-resistant healthcare-associated bacteria share MGEs across large phylogenetic distances^7–9^. Understanding the dynamics of MGE transfer in clinical settings can uncover important epidemiologic links that are not currently identified by traditional infection control methodologies^1,10,11^.

Methods to identify and track the movement of MGEs among bacterial populations on short timescales are limited. Bacterial whole-genome sequencing has transformed infectious disease epidemiology over the last decade^12^, providing powerful new tools to identify and intervene against outbreaks^13^. Despite these advances, efforts to track MGE movement have focused almost exclusively on drug resistance and virulence genes^5,7,11,14^, often ignoring the broader genomic context of the mobile elements themselves. Many studies rely on the identification of plasmid replicons, transposases, and other “marker genes”,^15^ an approach that oversimplifies the diversity of MGEs and may lead to incomplete or erroneous conclusions about their epidemiology. While querying databases containing curated MGE-associated sequences is useful for the rapid screening of clinical isolates for known MGEs, it will not capture novel MGEs. Additionally, whole-genome sequencing using short-read technologies generates genome assemblies that usually do not resolve MGE sequences, due to the abundance of repetitive elements that MGEs often contain^16^. Advances in long-read sequencing can mitigate this problem; the combination of short- and long-read sequence data can allow the genomic context of chromosomal and extrachromosomal MGEs to be precisely visualized^7,17,18^. Finally, studying the epidemiology of MGEs in clinical settings requires detailed individual-level patient clinical data, without which HGT occurrence in the hospital cannot be identified^18^.

Here we performed an alignment-based screen for MGEs in a large and diverse collection of bacterial genomes sampled from infections within a single hospital over an 18-month time period. We identified complete and fragmented MGEs that were identical in nucleotide sequence and occurred in the genomes of bacteria belonging to different genera. Because they are identical, we suspect that these MGEs have recently transferred between bacteria within the hospital setting. Further analysis using long-read sequencing and referenced-based resolution of distinct MGEs enabled us to precisely characterize MGE architecture and cargo, and to track MGE occurrence over time. Cross-referencing our results with available patient metadata allowed us to follow these elements as they emerged and were maintained among nosocomial bacterial populations.

## RESULTS

### Identification of MGEs shared across bacterial genera in a single hospital

Our experimental workflow is depicted in Fig. 1A. To identify genetic material being shared between distantly related bacteria in the hospital setting, we screened a dataset containing 2,173 whole-genome sequences of clinical isolates of high-priority Gram-positive and Gram-negative bacteria collected from a single hospital over an 18-month period as part of the Enhanced Detection System for Hospital-Acquired Transmission (EDS-HAT) project at the University of Pittsburgh^19^ (Methods). To have maximal contrast in our identification of MGEs, we focused on identical sequences found in the genomes of bacteria belonging to different genera. We performed an all-by-all alignment of the 2,173 genomes in the dataset using nucmer^20^, and filtered the results to retain alignments of at least 5kb that shared 100% identity between bacteria of different genera. The resulting sequences were extracted and clustered using Cytoscape (Fig. 1B). This approach identified putative MGE sequences in 196 genomes belonging to 11 genera, which could be grouped into 51 clusters of related MGEs. These MGE clusters ranged in size from two to 52 genomes, and comprised two, three, or four different genera (Fig. 1B). MGE sequences were found predominantly among Gram-negative *Enterobacteriaceae*, particularly *Klebsiella spp., Escherichia coli*, and *Citrobacter spp.* (Fig. 1C). Annotation of clustered sequences confirmed that more than 80% of the MGE clusters encoded one or more genes involved in DNA mobilization, plasmid replication, or another mobile function presumably involved in HGT (Fig. 1D). Somewhat surprisingly, only about one-quarter of the MGE clusters contained antimicrobial resistance genes, including genes encoding resistance to aminoglycosides, antifolates, beta-lactams, macrolides, quinolones, sulphonamides, and tetracyclines (Fig. 1D, 1E). Finally, 8/51 MGE clusters encoded genes and operons whose products were predicted to interact with metals, including arsenic, copper, mercury, nickel, and silver (Fig. 1D). Collectively, these results indicate that our unbiased, alignment-based method successfully identified putative MGEs, particularly in pathogens known to engage in HGT^2,21^.

**Figure 1.**
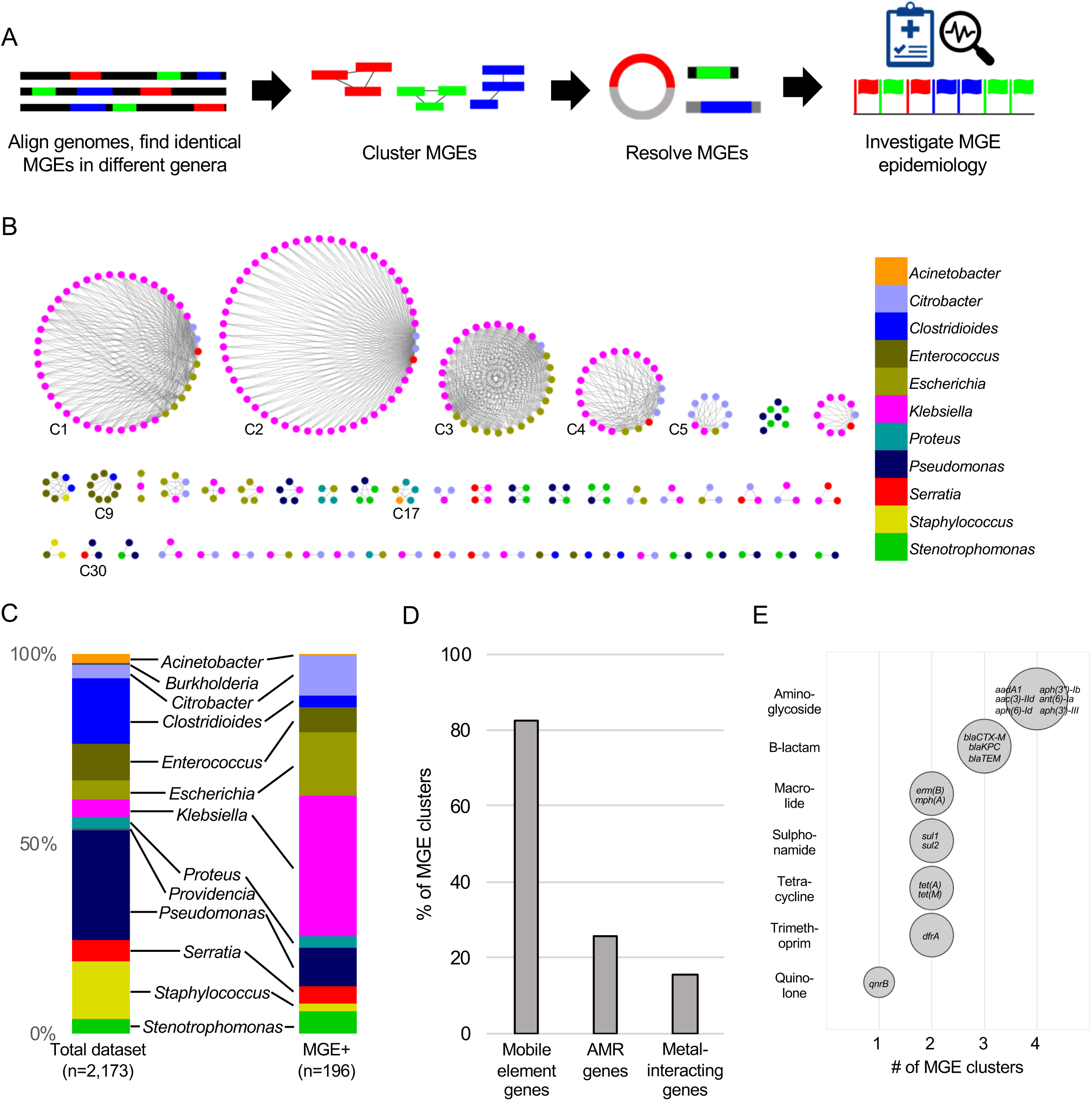
Identical mobile genetic elements (MGEs) shared across bacterial genera in a single hospital. (A) Approach to identify and characterize MGEs. (B) MGEs identified by the approach shown in (A). 51 MGE clusters found in distinct genera visualized with Cytoscape. Nodes represent bacterial isolates and are color-coded by genus. Edges connect nodes from different genera sharing >5kb of sequence at 100% nucleotide sequence identity. Clusters examined more closely in subsequent figures are labeled. (C) Genus distribution of all 2,173 genomes in the dataset (left) and the 196 isolates with MGEs shared across genera (right). (D) Prevalence of annotated mobile element, antimicrobial resistance (AMR) and metal-interacting genes among 51 MGE clusters. (E) Summary of AMR genes identified in MGE clusters. Genes are grouped by antibiotic class and bubble sizes correspond to prevalence among the MGE clusters shown in (B). AMR gene names are listed inside each bubble.

To assess the phylogenetic distribution of the putative MGEs we identified, we constructed a core gene phylogeny of the 196 genomes encoding one or more MGE clusters using the Genome Taxonomy Database Tool Kit (GTDBTK)^22^ (Fig. 2). MGE clusters were often found among bacteria in related genera, in particular the *Enterobacteriaceae.* We did not observe any MGEs that were present in both Gram-positive and Gram-negative isolate genomes, but we did find MGEs in the genomes of distantly related bacteria. For example, we identified an MGE carrying three aminoglycoside resistance genes that was identical in sequence between a vancomycin resistance-encoding plasmid carried by *E. faecium* and the *C. difficile* chromosome (MGE cluster C9, Fig. 3A). The *C. difficile* strain carrying this element was previously found to also harbor an *npmA* aminoglycoside resistance gene^23^. We also found portions of an integrative conjugative element that were identical between two *P. aeruginosa* isolates and a S. *marcescens* isolate (MGE cluster C30, Fig. 3B). Identical regions of this element included formaldehyde resistance genes and Uvr endonucleases. Finally, we detected complete and identical Tn7 transposons in the genomes of *A. baumannii, E. coli*, and *P. mirabilis* isolates (MGE cluster C17, Fig. 3C). The Tn7 sequence we detected was also identical to the Tn7 sequence of pR721, an *E. coli* plasmid that was first described in 1990 and was sequenced in 2014^24^. Taken together, these results indicate that while many of the sequences we identified were from MGEs shared between related bacterial genera, our approach also identified partial or complete MGEs that were identical in the genomes of distantly related pathogens.

**Figure 2.**
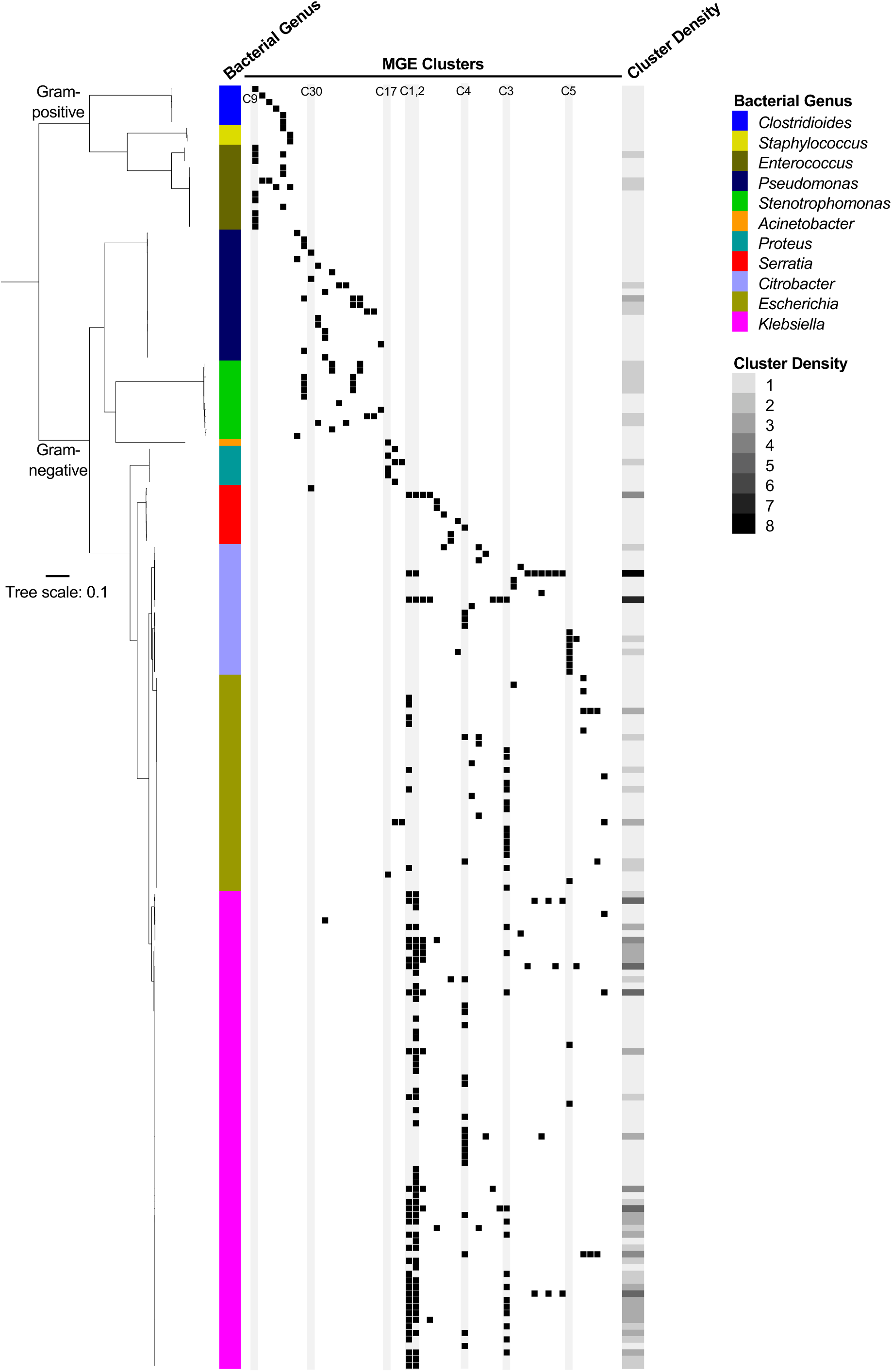
Phylogenetic distribution of MGE clusters across 196 genomes. Black squares mark the presence of one or more MGE clusters in each genome, with each column corresponding to a different MGE cluster. The heat map to the right shows MGE cluster density (i.e. total number of cross-genus MGEs) in each bacterial genome. Clusters examined more closely in subsequent figures are labeled and shaded in gray.

**Figure 3.**
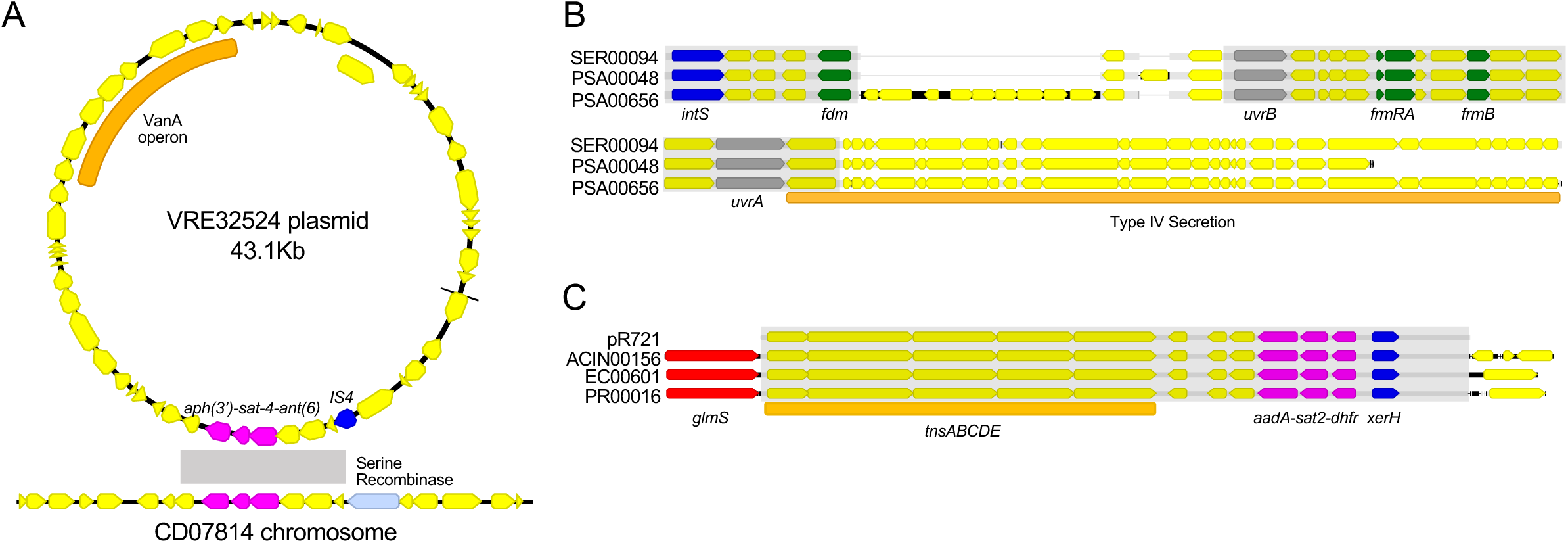
Examples of MGE sharing across genera. (A) Genes shared between a vancomycin-resistant *E. faecium* (VRE) plasmid and a *C. difficile* chromosome (MGE cluster C9). The VanA operon, conferring vancomycin resistance, is marked with an orange bar. Shared drug resistance genes are colored magenta, and mobile element genes are colored blue. Gray shading marks a stretch of DNA sequence that is 100% identical between isolates. (B) Identical portions of an integrated conjugative element (MGE cluster C30) shared between an *S. marcescens* genome (SER00094) and two *P. aeruginosa* genomes (PSA00048 and PSA00656). Blue = *intS* integrase; green = formaldehyde resistance genes; gray = UvrABC system genes. Type IV secretion machinery is marked with an orange bar, and gray shading marks sequences that are 100% identical between isolates. (C) Identical Tn7 transposons shared between *A. baumannii, E. coli*, and *P. mirabilis* (MGE cluster C17). The Tn7 sequence of the pR721 plasmid is shown at the top. The *tnsABCDE* transposon machinery is marked with an orange bar, and the *glmS* gene, which flanks the Tn7 insertion site, is colored red. Shared drug resistance genes are colored magenta, and an *xerH* tyrosine recombinase is colored blue. Gray shading marks sequences that are 100% identical between isolates.

### MGEs often reside on larger elements in different combinations and contexts

To further investigate the genomic context of the MGEs identified, we selected representative isolates from the largest MGE clusters for long-read sequencing using Oxford Nanopore technology. Hybrid assembly using short Illumina reads and long Nanopore reads generated highly contiguous chromosomal and plasmid sequences, which allowed us to resolve larger elements carrying one or more of the most prevalent MGE clusters (Table 1). We found that several of the smaller and more prevalent MGEs were carried on a variety of different plasmid and chromosomal elements, which we designated as “MGE lineages” (Table 1, Fig. 4A). These smaller MGEs co-occurred in different orders, orientations, and combinations on the larger elements. This kind of “nesting” of MGEs within larger mobile elements has been previously observed^6^, and our findings further support the mosaic, mix-and-match nature of the smaller MGEs we identified. We also confirmed that these MGEs were truly mobile, since they appeared to be able to move independently between multiple distinct larger mobile elements. A closer examination of the three largest MGE clusters (C1, C2, C3) showed that C1 sequences did not all share a common “core” nucleotide sequence, but rather could be aligned in a pairwise fashion to generate a contiguous sequence (Fig. 4B). MGE clusters C2 and C3, on the other hand, did contain “core” sequences that were present in all genomes carrying the MGE (Fig. 4C, 4D).

**Table 1.** MGE-encoding lineages and associated antibiotic resistance and metal interaction gene contents.

| MGE Lineage<sup>a</sup> | Length (kb) | Replicons (PlasmidFinder) | Antibiotic Resistance (ResFinder) | Metal Interaction (Prokka) |
| --- | --- | --- | --- | --- |
| cEC00609 | 39 | None | *aac(3)-IIa, aac(6')-Ib-cr, blaCTX-M-1, blaOXA-1, catB3, tet(A)* | None |
| pCB00017_2 | 196.8 | IncFIB(K), IncFII(K) | *aac(6')-Ib-cr, aph(3'')-Ib, aph(6)-Id, blaCTX-M-15, blaOXA-1, blaTEM-1B, catB3, qnrB1, tet(A), sul2* | *copD* operon, *pcoE, silE, silP, ars* operon |
| pCB00028_2 | 383.1 | IncHI2, IncHI2A, repA | *aac(3)-IIa, aac(6')-Ib-cr, aadA1, aph(3'')-Ib, aph(6)-Id, blaCTX-M-15, blaOXA-1, blaTEM-1B, catA1, catB3, dfrA14, sul2, tet(A)* | *pcoE, merR, merB* |
| pEC00668_2 | 145.4 | IncFIA, IncFIB(AP001918), IncFII(pRSB107) | *aac(6)-Id, aph(3'')-Ib, dfrA14, blaTEM-1B, mph(A), sul2* | *efeU, merA, merC, merP, merR, merT* |
| pEC00690_2 | 106.8 | IncFIA, IncFII | *aac(6')-Ibcr, blaOXA-1, catB3, tet(A)* | *efeU* |
| pKLP00149_2 | 165.2 | FII(pBK30683) | *aac(6')-Ib, aac(6')-Ib-cr, aadA1, aph(3'')-Ib, aph(6)-Id, blaKPC-3, blaOXA-9, blaSHV-182, blaTEM-1A, dfrA14, sul2* | *csoR* |
| pKLP00155_6 | 9.5 | ColRNAI | None | None |
| pKLP00161_2 | 236.5 | IncFIB(K), IncFII(K) | *aac(6')-Ib-cr, aph(3'')-Ib, aph(6)-Id, blaCTX-M-15, blaOXA-1, blaTEM-1B, dfrA14, qnrB1, sul2, tet(A)* | *copD* operon, *pcoC, pcoE, silE, silP, ars* operon |
| pKLP00177_3 | 170.8 | IncFIB(K) | *aac(3)-IIa, aac(6')-Ib-cr, aph(3'')-Ib, aph(6)-Id, blaCTX-M-15, blaOXA-1, blaTEM-1B, catB3, dfrA14, qnrB1, sul2, tet(A)* | *copD* operon, *pcoC, pcoE, silE, silP, ars* operon |
| pKLP00182_3 | 15.8 | None | *aac(6')-Ib-cr, blaOXA-1, catB3, dfrA14, tet(A)* | None |
| pKLP00215_4 | 113.6 | IncFIB(pQil), IncFII(K) | *blaKPC-2, blaOXA-9, blaTEM-1A* | *merB, merR* |
| pKLP00218_2 | 164.7 | IncFIB(K), IncFII(K) | *aph(3'')-Ib, aph(6)-Id, blaCTX-M-15, blaTEM-1B, dfrA14, sul2* | *copD* operon, *pcoC, pcoE, silE, silP, ars* operon |
| pKLP00221_2 | 242.3 | ColRNAI, IncFIB(K), IncFII(K) | *aac(6')-Ib, aada2, aph(3')-1a, blaKPC-2, blaOXA-9, blaTEM-1A, catA1, dfrA12, mph(A), sul1* | *copD* operon, *pcoC, pcoE, silE, silP, ars* operon |
| pKLO00017_2 | 226.2 | IncFIB(K), IncFII(K) | *aph(3'')-Ib, aph(6)-Id, blaCTX-M-15, blaTEM-1B, dfrA14, sul2* | *copD* operon, *pcoC, pcoE, silE, silP, ars* operon |

<sup>a</sup>MGE lineage names include location (c = chromosome, p = plasmid), name of the reference isolate sequenced, and assembly contig number (\_2, \_3, \_4, \_6).

**Figure 4.**
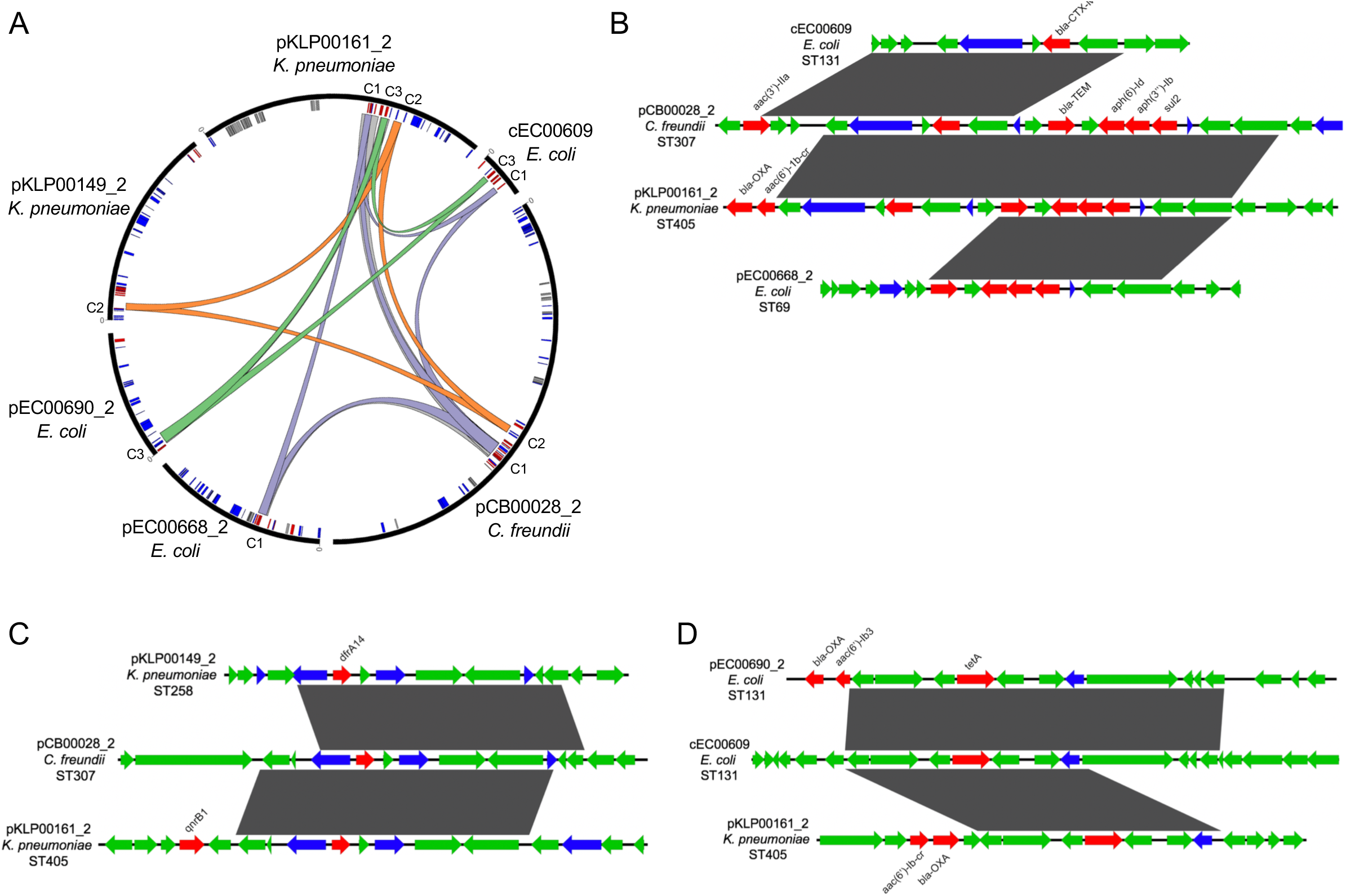
Mosaicism of MGE clusters present on diverse elements. (A) Circular plot of six distinct sequence elements (black bars) that encode MGE clusters C1, C2, and C3. Lowercase letters in sequence names indicate element type (c = chromosome, p = plasmid). Homologous cluster sequences are connected to one another with colored links (purple = C1, orange = C2, green = C3, gray = other). Inner circle depicts MGE genes involved in MGE mobilization (blue), antibiotic resistance (red) and metal interaction (gray). (B-D) Alignments of sequences grouped into MGE clusters C1 (B), C2 (C), and C3 (D) from the larger MGEs displayed in (A). ORFs are colored by function (blue = mobilization, red = antibiotic resistance, green = other/hypothetical). Antibiotic resistance genes are labeled above and dark gray blocks connect sequences that are identical over at least 5kb.

### Plasmids carrying MGE clusters are found in multiple sequence types, species, and genera circulating in the same hospital

More than half (104/196) of the MGE-carrying genomes in our dataset contained one or more of the five most prevalent MGEs we identified (C1-C5, Fig. 1B). All five MGEs were small (usually less than 10kb), and were predicted to be carried on plasmids shared between *Enterobacteriaceae*. We set out to resolve the genomic context of each of these five MGEs in all isolates containing them. We used an iterative approach involving long-read sequencing and hybrid assembly of representative isolates to generate reference sequences of MGE-containing elements (chromosomal or plasmid), followed by mapping of contigs from Illumina-only assemblies to these reference sequences to assess their coverage in every genome (Methods). This approach allowed us to query the presence of plasmids and chromosomal elements from genomes sequenced with Ilumina technology alone, without requiring long-read sequencing of all isolates or relying on external reference sequences. We found that 11 of the 104 isolates (all *E. coli*) carried cluster C1 and C3 MGEs on their chromosome, while the remaining 93 isolates carried clustered MGEs on 17 distinct plasmids. Seven of these plasmids were present in only one isolate in the dataset, but 10 plasmids appeared to be shared between more than one isolate (Table 1, Fig. 5). We also conducted the same reference-based coverage analysis for all 2,173 genomes in the original dataset, and identified an additional 16 isolates with >90% coverage of an MGE lineage.

**Figure 5:**
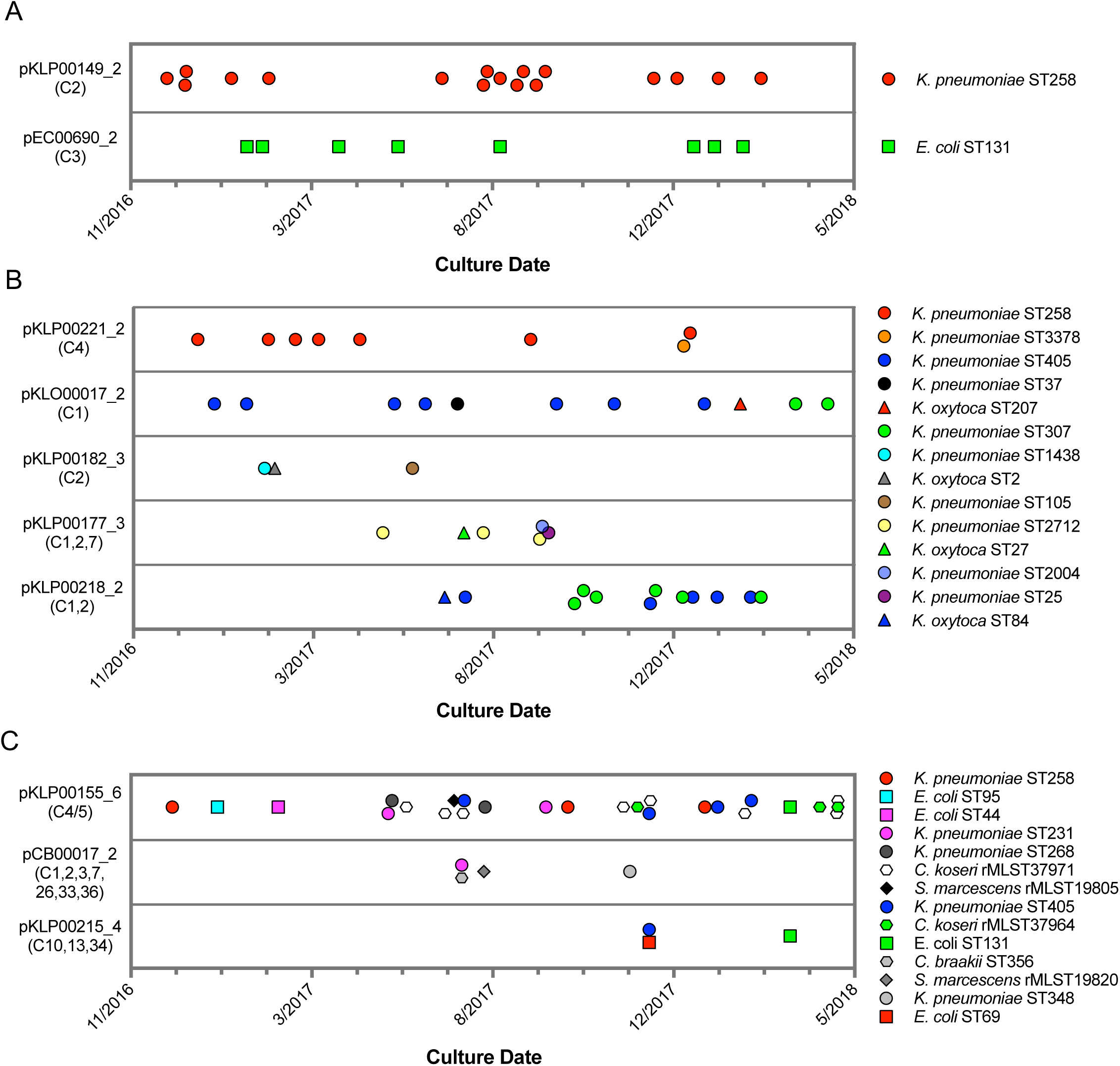
Timelines of plasmid lineage occurrence among isolates of the same ST (A), same genus (B), or different genera (C). Timelines show the culture date of isolates predicted to encode plasmids belonging to the same lineage, based on coverage mapping to reference plasmids listed to the left of each timeline. The MGE clusters carried by each plasmid are listed in parentheses below the plasmid name. More information about each plasmid is provided in Table 1. Shape and color of data points correspond to bacterial species and ST, respectively.

While all of the MGEs we originally identified were present in the genomes of bacteria belonging to different genera, the plasmids that we resolved were variable in how widely they were shared. For example, some plasmids were only found among isolates belonging to a single species and multilocus sequence type (ST), suggesting that they were likely transmitted between patients along with the bacteria that were carrying them (Fig. 5A). These included a *blaKPC-3* carbapenemase-encoding plasmid (pKLP00149_2) found in *K. pneumoniae* isolates belonging to ST258, a multidrug-resistant and highly virulent hospital-adapted bacterial lineage that has recently undergone clonal expansion in our hospital^18^. We also found a *blaOXA-1* extended spectrum beta-lactamase-encoding plasmid in *E. coli* isolates belonging to ST131, another multidrug-resistant and hypervirulent bacterial lineage^25^. In addition to plasmids that occurred in bacteria belonging to the same ST, we also identified plasmids that were present in isolates belonging to different STs of the same species, or in different species of the same genus (Fig. 5B). All isolates in this case were *K. pneumoniae* or *K. oxytoca*, suggesting widespread sharing of plasmids between distinct *Klebsiella* species and STs. The plasmids often carried antibiotic resistance genes, and many also carried metal interaction genes (Table 1). Finally, we identified three different plasmids that were shared between different bacterial genera all belonging to the *Enterobacteriaceae* (Fig. 5C). One small plasmid (pKLP00155_6) carrying the colicin bacterial toxin was found in 26 isolates belonging to 10 different STs and four different genera. Taken together, these results indicate that some plasmids carrying putative MGEs were likely inherited vertically as bacteria were transmitted between patients in the hospital, while others appear to have transferred independently of bacterial transmission.

### Likely HGT across genera within individual patients

By cross-referencing the isolates containing MGE sequences with de-identified patient data, we found two instances where identical MGEs were found in pairs of isolates of different genera that were collected from the same patient, on the same date, and from the same sample source. To resolve the complete MGE profiles of these cases, we performed long-read sequencing and hybrid assembly on all genomes involved (Fig. 6). A *K. pneumoniae* ST405 isolate (KLP00215) and an *E. coli* ST69 isolate (EC00678) collected from a tissue infection from Patient A each harbored a 113.6kb IncFIB(pQil)/IncFII(K) plasmid carrying *blaKPC, blaTEM*, and *blaOXA* enzymes, as well as a mercury detoxification operon (Fig. 6A, B). In addition, an isolate from a second patient (Patient B, EC00701, *E. coli* ST131), also encoded a nearly identical plasmid. A systematic chart review for Patients A and B revealed that they occupied adjacent hospital rooms for four days during a time period after Patient A’s isolates were collected but before Patient B’s isolate was collected. During this time the two patients would have shared the same healthcare staff, who might have transferred bacteria between them.

**Figure 6:**
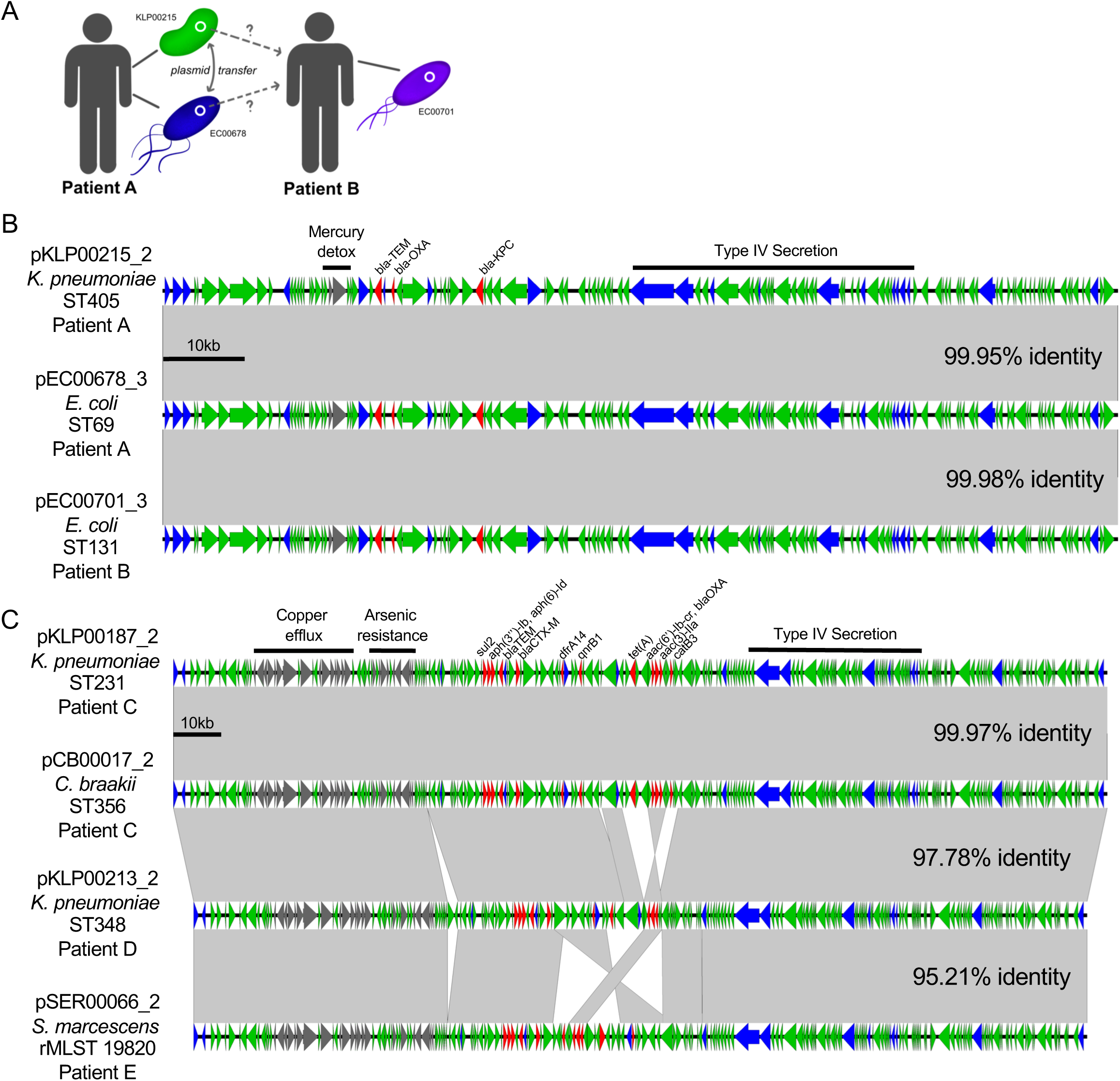
Cross-genus transfer of plasmids within and between patients. (A) Schematic diagram showing *K. pneumoniae* and *E. coli* isolates bearing the same plasmid sampled from two patients. (B) Nucleotide alignment of the plasmid presumably transferred within and between the patients shown in (A). A 113.6kb IncFIB(pQil)/IncFII(K) carbapenemase-encoding plasmid was resolved from two genomes of different bacterial isolates from the same clinical specimen from Patient A. A nearly identical plasmid was also identified in an isolate from Patient B, who occupied a hospital room adjacent to Patient A. (C) Alignment of a 196.8kb IncFIB(K)/IncFII(K) multidrug-resistance plasmid resolved from two genomes of different bacterial isolates from the same clinical specimen from Patient C. Similar plasmids were also found in isolates from two additional patients (Patient D and Patient E), who had no identifiable epidemiologic links with Patient C. ORFs are colored by function (blue = mobilization, red = antibiotic resistance, gray = metal-interacting, green = other/hypothetical). Antibiotic resistance genes, metal-interacting operons, and Type IV secretion components are labeled. Shading between sequences indicates regions >5kb with >99.9% identity, and pairwise identities across the entire plasmid are noted to the right.

In the second case of putative within-patient HGT, a *K. pneumoniae* ST231 isolate (KLP00187) and a *C. braakii* ST356 isolate (CB00017) were both collected from the same urine sample of Patient C (Fig. 6B). Both isolates carried nearly identical 196.8kb IncFIB(K)/IncFII(K) plasmids conferring resistance to aminoglycosides, beta-lactams, chloramphenicol, fluoroquinolones, sulfonamides, tetracyclines, and trimethoprim, as well as operons encoding copper and arsenic resistance. In addition, isolates from two subsequent patients (Patient D and Patient E) also carried plasmids belonging to the same lineage as the plasmid shared between KLP00187 and CB00017. Alignment of the sequences of all four plasmids showed that the plasmids isolated from Patient C were nearly identical, while the plasmids from Patients D and E had small differences in their gene content and organization (Fig. 6B). Systematic chart review did not identify any strong epidemiologic links between the three patients, suggesting that this plasmid was not passed directly between these patients and might instead have transferred via additional bacterial isolates or populations that were not sampled.

## DISCUSSION

In this study, we identified MGEs in a large dataset of whole-genome sequences of clinical bacterial isolates collected over an 18-month period from a single hospital. We identified, clustered, and characterized identical sequences found in multiple distinct genera, and in the process uncovered both expected and unexpected cases of MGE occurrence. We confirmed that some of the most common MGEs identified were fragments of larger mobile elements. We performed long-read sequencing to resolve these larger elements, which were almost always plasmids. When we traced the presence of various plasmid lineages over time, we found some that were likely transmitted vertically along with the bacteria carrying them, and others that appeared to be transferred horizontally between unrelated bacteria.

Our study adds to the body of knowledge of HGT in hospital settings in new and important ways. We analyzed a large dataset of clinical isolates collected from a single health system, and used a systematic and unbiased approach to identify MGEs regardless of their type or gene content. While prior studies have used genomic epidemiology to study how HGT contributes to the transmission, persistence, and virulence of bacterial pathogens^4,5,19,20^, the technical challenges of resolving MGEs from whole-genome sequencing data have limited the scope of these findings^16^. Other studies have deliberately tracked HGT in healthcare settings by focusing either on mobile genes of interest, such as those encoding drug resistance^7,9,14^, or on specific classes of MGEs^28^. Both of these approaches can generate biased interpretations of the driving forces behind HGT in clinical settings. For this reason we selected a pairwise alignment-based approach, whereby we only looked for identical sequences in the genomes of very distantly related bacteria. In doing so, we did not limit ourselves to only looking for “known” MGEs, and thus obtained a more accurate and comprehensive overview of the dynamics of HGT between bacterial genera in our hospital.

What might cause horizontally-transferred nucleotide sequences to be found at very high identity within phylogenetically distinct bacteria? We predicted that there might be two possible causes: Either the sequences we identified represent MGEs that recently underwent HGT and have not had time to diverge from one another, or they represent genetic elements that are highly intolerant to mutation. We suspect that our dataset contains both cases. In the two instances of likely within-patient HGT, both plasmids isolated from the same patient were nearly identical to one another, suggesting that they were indeed transferred shortly before the bacteria were isolated. In both cases we also observed similar plasmids in the genomes of isolates from other patients, but we identified a likely route of transfer between patients only in the case where the subsequent plasmid was also nearly identical. This finding further supports the idea that high plasmid identity is evidence of recent transfer. On the other hand, the Tn7 transposon sequence we uncovered that was identical in bacterial isolates from three different genera was also identical to over two dozen publicly available genome sequences queried through a standard NCBI BLAST search. The insertion of the Tn7 transposon downstream of *glmS* in all of our isolates suggests TnsD-mediated transposition^29^, but the reason why the entire transposon sequence is so highly conserved is unclear.

The vast majority of MGE sequences identified through our approach contained signatures of mobile elements, and our follow-up work demonstrated that they could very likely move independently and assemble mosaically on larger mobile elements, such as plasmids, integrative conjugative elements, and other genomic islands. Antibiotic resistance genes were present in fewer MGE clusters than we anticipated, given how many resistance genes are known to be MGE-associated. Our follow-up analysis showed, however, that resistance genes were indeed highly prevalent among the larger MGEs that we resolved. This suggests that resistance genes often reside on smaller and more variable elements, which would have been filtered out by the parameters of our initial screen. A recent study of clinical *K. pneumoniae* genomes showed that while antibiotic resistance genes were largely maintained at the population level, they were variably present on different MGEs that fluctuated in their prevalence over time^24^. Finally, we were somewhat surprised by the large number of metal-interacting genes and operons within the MGEs that we identified. While metal-interacting genes and operons have been hypothesized to confer disinfectant tolerance and increased virulence^30,31^, precisely how these elements might increase bacterial survival in the hospital environment and/or contribute to infection requires further study.

Identification of risk factors and common exposures for HGT has previously been proposed^1,14,18,32^, but the results of prior efforts have been limited because large genomic datasets from single health systems with corresponding epidemiologic data have not been widely available^33^. The use of routine whole-genome sequencing for outbreak surveillance in our hospital has allowed us to begin to study how the transmission of MGEs might be similar or different from bacterial transmission. In addition to finding evidence of vertical transfer of plasmids accompanying bacterial transmission, we also identified several cases in which the same MGE lineage was identified in two or more isolates of different sequence types, species, or genera. In some cases, these isolates were collected within days or weeks of one another. This finding underscores how rapidly MGEs can move between bacterial populations, particularly in hospitalized patients^1,21^, and highlights the importance of pairing genome sequencing with epidemiologic data to uncover routes of MGE transmission.

There were several limitations to our study. First, the dataset that we used only contained genomes of isolates from clinical infections from a pre-selected list of species, and did not include environmental samples or isolates from patient colonization. Second, our method to screen for putative MGE sequences based on cross-genus alignment was based on somewhat arbitrary cutoffs, and we largely ignored MGEs that only transferred between bacteria within a single genus. Additionally, the cross-genus parameter we employed may have artificially enriched the number of MGEs we identified among *Enterobacteriaceae*, which are known to readily undergo HGT with one another^7^. Third, we assigned MGE lineages relative to single reference sequences and based on our analysis on reference sequence coverage; subsequent MGEs that either gained additional sequence or rearranged their contents would still be assigned to the same lineage, even though they may have diverged substantially from the reference MGE^6^. Finally, this study was based exclusively on comparative genome analyses, and the MGEs we resolved from clinical isolate genomes were not queried for their capacity to undergo HGT *in vitro*.

In conclusion, we have shown how bacterial whole genome sequence data, which is increasingly being generated in clinical settings, can be leveraged to study the dynamics of HGT between drug-resistant bacterial pathogens within a single hospital. Our future work will include further characterization of the MGEs we resolved, assessment of MGE sharing across closer genetic distances, and incorporation of additional epidemiologic information to identify shared exposures and possible routes for MGE transfer independent from bacterial transmission. Ultimately we aim to develop this analysis into a reliable method that can generate actionable information and enhance traditional approaches to prevent and control multidrug-resistant bacterial infections.

## METHODS

### Isolate collection

Isolates were collected through the Enhanced Detection System for Hospital-Acquired Transmission (EDS-HAT) project at the University of Pittsburgh^19^. Eligibility of bacterial isolates for genome sequencing under EDS-HAT required positive clinical culture for high-priority and multidrug-resistant bacterial pathogens with either of the following criteria: >3 hospital days after admission, and/or any procedure or prior inpatient stay in the 30 days prior to isolate collection. Pathogens collected included: *Acinetobacter spp., Burkholderia spp., Citrobacter spp., Clostridioides difficile*, vancomycin-resistant *Enterococcus spp.*, extended-spectrum beta-lactamase (ESBL)-producing *E. coli*, ESBL-producing *Klebsiella spp., Proteus spp., Providencia spp., Pseudomonas spp., Serratia spp., Stenotrophomonas spp.*, and methicillin-resistant *S. aureus*. Eligible isolates were identified using TheraDoc software (Version 4.6, Premier, Inc., Charlotte, NC). The EDS-HAT project involves no contact with human subjects; the project was approved by the University of Pittsburgh Institutional Review Board and was classified as being exempt from informed consent. De-identified patient IDs and culture dates were utilized in downstream analysis.

### Whole genome sequencing and analysis

Genomic DNA was extracted from pure overnight cultures of single bacterial colonies using a Qiagen DNeasy Tissue Kit according to manufacturer’s instructions (Qiagen, Germantown, MD). Illumina library construction and sequencing were conducted using the Illumina Nextera DNA Sample Prep Kit with 150bp paired-end reads, and libraries were sequenced on the NextSeq sequencing platform (Illumina, San Diego, CA). Selected isolates were also sequenced with long-read technology on a MinION device (Oxford Nanopore Technologies, Oxford, United Kingdom). Long-read sequencing libraries were prepared and multiplexed using a rapid multiplex barcoding kit (catalog RBK-004), and were sequenced on R9.4.1 flow cells. Base-calling on raw reads was performed using Albacore v2.3.3 or Guppy v2.3.1 (Oxford Nanopore Technologies, Oxford, United Kingdom).

Illumina sequencing data were processed with Trim Galore v0.6.1 to remove sequencing adaptors, low-quality bases, and poor-quality reads. Bacterial species were assigned by k-mer clustering with Kraken v1.0^34^ and RefSeq^35^ databases. Genomes were assembled with SPAdes v3.11^36^, and assembly quality was verified using QUAST^37^. Genomes were annotated with Prokka v1.13^38^. Multi-locus sequence types (STs) were assigned using PubMLST typing schemes with mlst v2.16.1^39,40^, and ribosomal sequence types (rMLSTs) for isolates not assigned an ST were approximated by alignment to rMLST reference sequences. Long-read sequence data was combined with Illumina data for the same isolate, and hybrid assembly was conducted using Unicycler v0.4.7 or v0.4.8-beta^41^.

### Identification and phylogenetic analysis of putative MGEs

Illumina genome assemblies were screened all-by-all against one another to identify alignments of at least 5,000bp at 100% identity using nucmer v4.0.0beta2^20^. The nucmer output was filtered to only include alignments between bacterial isolates of different genera. Nucleotide sequences from the resulting alignments were then extracted and compared against one another by all-by-all BLASTn v2.7.1^42^. Results were filtered to only include nucleotide sequences having 100% identity over at least 5,000bp to at least one sequence from another genus. The resulting comparisons were clustered and visualized using Cytoscape v3.7.1^43^. A phylogeny of MGE-encoding genomes was constructed using the Genome Taxonomy Database Tool Kit (GTDBTK)^22^. Briefly, translated amino acid sequences of 120 ubiquitous bacterial genes were generated, concatenated, and aligned using GTDBTK’s *identify* pipeline. The resulting multiple sequence alignment was masked for gaps and uncertainties, then a phylogenetic tree was generated using RAxML v8.0.26 with the PROTGAMMA substitution model^44^ and 1000 bootstraps.

### Characterization of MGE fragments and assignment of chromosomal element and plasmid lineages

The longest nucleotide sequence in each MGE cluster was considered representative of that cluster, and was annotated with Prokka v1.13. Representative sequences were compared to publicly available genomes by BLASTn v2.7.1 against the NCBI Nucleotide database. Antibiotic resistance genes were identified by a BLASTn-based search against the CARD v3.0.1^29^ and ResFinder v3.2^46^ databases, and plasmid replicons were identified by a BLASTn-based search against the PlasmidFinder database v2.0.2^47^. Additional features of each MGE cluster were identified by consulting annotations assigned by Prokka. MGEs were aligned to one another using Geneious v11.1.5 (Biomatters Ltd., Aukland, New Zealand) and EasyFig v2.2.2^48^.

To resolve larger mobile elements encodings MGEs C1-C5, we first selected the earliest isolate containing each MGE for long-read sequencing and hybrid assembly. The closed, MGE-encoding element (plasmid or chromosomal) from this earliest isolate was used as a reference for mapping contigs from Illumina assemblies from all other isolates using BLASTn. Briefly, contigs from Illumina-only assemblies were aligned to each reference MGE-encoding element, and isolates having at least 90% coverage of the reference element were assigned to that element’s “lineage.” Among isolates having less than 90% coverage, a representative was again selected for long-read sequencing and hybrid assembly, and the process was repeated until all 104 isolates had been assigned to a lineage. Lineages were named based on the MGE-containing element type (c = chromosomal, p = plasmid), the reference isolate, and the hybrid assembly contig number, denoted with an underscore at the end of the name. MGE cluster-containing plasmids resolved through hybrid assembly were also used as reference sequences to query their presence in the entire 2,173 genome data set using the same BLASTn coverage-based analysis as above. When isolate genomes showed high coverage of multiple reference plasmids, the longest plasmid having at least 90% coverage was recorded.

### Systematic chart review to assess epidemiologic links between patients with the same plasmids

Patients whose isolates carried the two plasmids found to putatively transfer within individual patients were reviewed using a systematic approach modified from previously published methodologies examining patient locations and procedures for potential similarities^49,50^. Patients were considered infected/colonized with the recovered plasmid on the day of the patients’ culture and all subsequent days. Potential transfer events were considered significant for locations if an uninfected/uncolonized patient was housed on the same unit location or service line location (units with shared staff) at the same time or different time as a patient infected/colonized with the plasmid, using a 60-day window prior to the newly infected/colonized patient’s culture date. Additionally, procedures (e.g. operation room procedures, bedside invasive procedures) were evaluated for commonalities among all patients 60 days prior to infection/colonization, as well as potential procedures contaminated by prior infected/colonized patients that could have transferred to newly infected/colonized patients, again using a 60-day window prior to the culture date. Procedures were deemed significant if >1 patient had a similar procedure, or if there was a shared procedure within the 60-day window.

## Acknowledgements

We gratefully acknowledge Chinelo Ezeonwuka, Kathleen Shutt, Daniel Snyder, Jieshi Chen, and Hayley Nordstrom for their helpful contributions to this study. This work was supported by a grant from the Competitive Medical Research Fund of the UPMC Health System to D.V.T., by NIAID grants R21Al109459 and R01AI127472 to L.H.H., and by the Department of Medicine at the University of Pittsburgh School of Medicine. The funders had no role in study design, data collection and interpretation, or the decision to submit the work for publication.

